# Human-derived alleles in *SOST* and *RUNX2* 3′UTRs cause differential regulation in a bone cell-line model

**DOI:** 10.1101/2021.04.21.440797

**Authors:** Juan Moriano, Núria Martínez-Gil, Alejandro Andirkó, Susana Balcells, Daniel Grinberg, Cedric Boeckx

## Abstract

The inquiry into the phenotypic features that set apart human species, such as a light, gracile skeleton and a rounded skull characteristic of *Homo sapiens*, can now benefit from the examination of ancient genomes. These have added a new layer of analysis allowing the identification of genetic differences between species like ours and our closest extinct relatives. Most of these genetic differences are non-coding changes with unknown functional consequences, and dissecting their putative regulatory effect remains challenging. Here we focus on the three prime untranslated regions (3’UTR), known to play a critical role in messenger RNA regulation and a plausible locus for divergent regulation between *Homo* species. We report a set of genes with derived 3’UTR changes in either the *Homo sapiens* or the Neanderthal/Denisovan lineages and experimentally evaluate the impact of 3’UTR variants in four genes: *E2F6*, *GLI3*, *RUNX2* and *SOST*. We performed a luciferase reporter assay in a bone cell-line model and found a statistically significant difference for the 3’UTR variants of *SOST* (*Homo sapiens*-derived) and *RUNX2* (Neanderthal/Denisovan-derived). The differential expression caused by these variants in our experimental model points to species differences in bone mineral density. Thus, this study adds insights into the functional effects of regulatory variants that emerged in recent human evolution.

## 1 Introduction

Among the features that distinguish *Homo sapiens* relative to extinct species of humans, such as Neanderthals, is a lighter bone structure. Neanderthals are known to have an overall higher bone density, probably coupled with higher muscle mass relative to *Homo sapiens* [1]. Different developmental trajectories [2] lead to clear differences in structures like the lower thorax [3], the lumbo-pelvic complex [3, 4], and in general give rise to relatively large limb length and a shorter stature [5].

While these morphological changes are linked to genetic variation, the underlying biological processes that drove the skeletal morphology of Neanderthals to a more robust shape are currently unknown. Owing to the appearance of the first high-quality Neanderthal/Denisovan genomes [6–8], the conventional approach based on the identification of missense changes can now be extended to the study of non-coding variants. Thus, efforts are needed to characterize the impact of non-coding variation at different layers of regulation and to infer the resulting phenotypic differences. Indeed, studies on human ancient genomes over the past decade have revealed that only a minor fraction (less than one hundred) of the genetic variants are fixed missense changes [9, 10].

Several lines of research have focused on single-nucleotide variants (SNVs) altering transcription factor binding sites [11–13], introgressed extinct human variants, deserts of introgression and positively selected regions that may underlie differential gene expression regulation in *Homo sapiens* when compared to the Neanderthal/Denisovan lineage (reviewed in [14]). The paleogenomic data have also been fruitfully exploited to identify ancient methylation maps. After pioneering work on the inference of epigenetic signals from ancient DNA [15–17], larger scale comparative analyses have allowed: i) the identification of differentially methylated regions predicted to impart regulation on certain anatomical structures, being the larynx and the protrusion of the face the more salient features assigned to the *Homo sapiens* lineage [18, 19], and ii) even the attempt to reconstruct the general skeletal anatomy of the Denisovans [20]. Precisely, the morphological features that characterize the modern human face—smaller brow ridges and nasal projections, reduced prognathism [21]—are thought to have emerged from a mild neurocristopathy underlying a ‘self-domestication’ process that led, as it is hypothesized, to unique aspects of *Homo sapiens* cognition and behavior [22].

To date, the experimental elucidation of the impact of all non-coding mutations that distinguish our species from our closest extinct relatives seems insurmountable. Substantial progress has been achieved through massive parallel reporter assays on specific cell-lines in the search of *Homo sapiens*-specific regulatory region activities [14, 23]. Here, we restrict our scope to a set of regulatory variants and to a bone cell-line model. On the one hand, bone cell-lines (e.g. osteoblasts) have been a useful model to inquire the evolutionary divergence in gene regulation and to infer its predicted effect on the evolution of anatomical structures (e.g. [17, 23]). On the other hand, the regulatory variants subject to scrutiny in this study belong to the 3’UTRs. These non-coding regions, located after the stop codon of messenger RNAs, are a site of regulation of protein expression and localization, containing binding sites for regulatory molecules such as microRNAs [24]. Evidence gathered from high-throughput sequencing studies converge on the study of 3’UTRs as a promising direction to investigate the molecular causes of disorders with developmental origin [24]. With a focus on recent human evolution, a miRNA containing a *Homo sapiens*-derived allele has been reported to underlie the differential regulation of two teeth formation-related genes in a reporter expression construct assay [25].

In the present study, we make use of paleogenomic data to identify genes whose 3’UTR acquired derived mutations in the *Homo sapiens* lineage respect to the Neanderthals/Denisovans (where the ancestral allelic version is found), and vice versa. An enrichment analysis reflects a diversity of gene ontology category terms, among which we find some related to anatomical features (limb development), dendritic spines or Wnt signaling pathway specifically for genes with *Homo sapiens*-derived changes. We selected four genes to experimentally test the impact of the 3’UTR species-specificvvariants: Three of them, *GLI3*, *RUNX2* and *SOST*, directly implicated in skeletal development and *E2F6*, a gene with comparatively lesser studied role in bone formation but whose 3’UTR overlaps with a putative positively-selected region [26]. We performed a luciferase reporter assay and found statistically significant differences for the 3’UTR regulatory variants of *SOST* and *RUNX2*. Particularly, we observed a downregulation of *SOST* and an upregulation of *RUNX2* caused by the Neanderthal/Denisovan non-coding variants, pointing to species-specific differences in bone mineral density (BMD) as the result of the differential expression of these two anabolic bone regulatory proteins.

## 2 Results

We processed a catalog of high-frequency SNVs distinguishing *Homo sapiens* and the Neanderthals and the Denisovans (from [10]) and integrated it with 3’UTR annotations from the Ensembl human assembly GRCh37 [27]. Out of the total 17268 genes with annotated 3’UTRs, 890 genes possess 3’UTRs with derived changes in our species, among which 64% (573 genes) have 3’UTR that did not accumulate derived changes in the homolog regions in the Neanderthal/Denisovan lineage. Complementarily, 2407 genes have Neanderthal/Denisovan-derived 3’UTR SNVs (87%, 2100 genes, with 3’UTRs depleted of *Homo sapiens*-derived changes). The complete catalog of genes with species-specific 3’UTR SNVs can be found in Supplementary Table 1. Enrichment analyses via the gprofiler2 R package [28] were performed to functionally evaluate the genes carrying species-specific changes in their 3’UTRs. General GO terms appear for terms shared between the datasets containing the genes with derived changes in either our species or the Neanderthals/Denisovans (890 and 2407 genes, respectively), among which we find the cerebral cortex, caudate and cerebellum structures (see Suppl. Table 1). It is noteworthy here that evidence from endocast landmark studies hints at differences in the morphology of the neurocranium of Neanderthals relative to extant humans, remarkably in the size and growth of the cerebellum and parietal regions (with a more elongated base of the neurocranium in the former species) [29–31]. Exclusive terms assigned to the *Homo sapiens* lineage are limb development (where differences between *Homo* species have been reported [5]), Wnt-signaling pathway (in relation to bone, particularly overrepresented in association studies of BMD [32, 33]), dendritic spine development, vesicle-mediated transport in synapse, along with several microRNAs and transcription factors (see Figure 1 and Suppl. Table 1). Assessing genes exclusively assigned to the *Homo sapiens* lineage (573 genes), however, returns few, unspecific GO terms.

**Figure 1:**
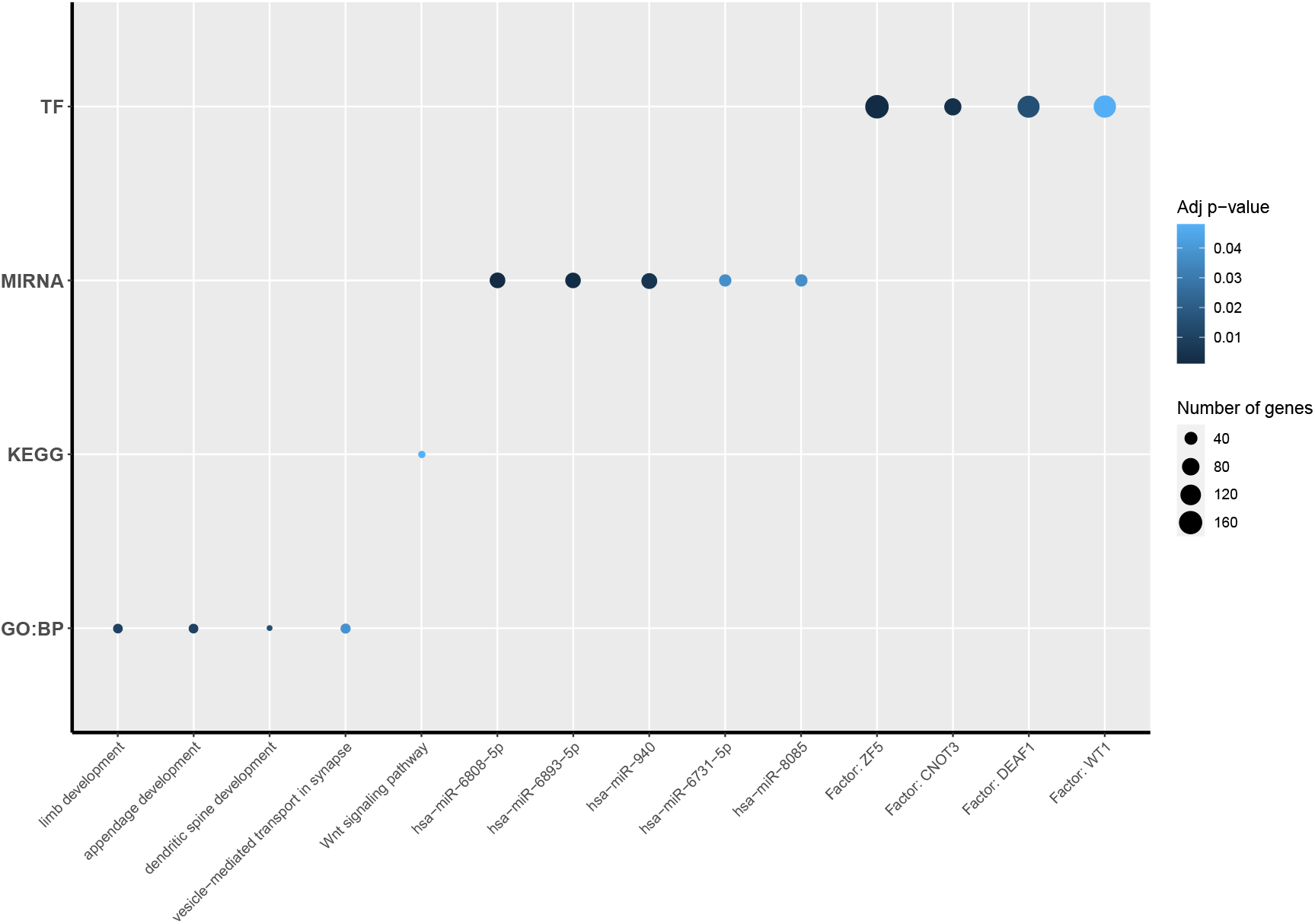
Exclusive GO terms assigned to genes with *Homo sapiens*-derived SNVs in their 3’UTRs. GO terms were filtered to select only those that are assigned to genes with *Homo sapiens*-derived 3’UTR SNVs and do not appear in the enrichment results for genes with Neanderthal/Denisovan-derived 3’UTR SNVs. Significant results were considered if *p <* 0.05. TF: Transcription factor; MIRNA: microRNAs; KEGG: Biological pathways; GO:BP: Biological process.

Next, we selected four candidate genes for experimental testing in a bone cell-line model (Saos-2 osteosarcoma cell-line). Among the genes with derived changes in only one of the two human lineages, we found three well-studied genes implicated in bone formation and homeostasis with pronounced phenotypic effects when altered: *GLI3*, critically involved in limb development and whose mutations have been identified in patients with polydactyly and macrocephaly [34]; *RUNX2*, a key transcription factor regulating osteoblast and chondrocyte differentiation, mutated in patients with cleidocranial dysplasia [35]; and *SOST*, an inhibitor of the canonical Wnt pathway that inhibits bone formation and that has been linked to sclerosteosis, van Buchem disease and craniodiaphyseal dysplasia [36, 37]. In addition, we also decided to evaluate the 3’UTR *Homo sapiens*-derived change of *E2F6*, a gene with little studied roles in skeletal development [38, 39], but whose 3’UTR overlaps a putative positively-selected region [26] and that also accumulated changes in the Neanderthal/Denisovan lineage. The *Homo sapiens*-derived change of *GLI3* also falls within a putative positively-selected region [26] (see Table 1 for a summary).

**Table 1:** Species-specific changes in 3’UTR. *GLI3* and *SOST* 3’UTRs contain *Homo sapiens*-derived SNVs and are depleted of Neanderthal/Denisovan changes. Inversely, the *RUNX2* 3’UTR contains only Neanderthal-derived SNVs. The *E2F6* 3’UTR contains derived changes in both lineages. *GLI3* and *E2F6 Homo sapiens*-specific 3’UTR variants overlap a putative positively-selected region [26]. Predicted microRNAs were assessed via the miRSNP online database http://bioinfo.bjmu.edu.cn/mirsnp/search/ HS: *Homo sapiens*; Nean/Den: Neanderthal/Denisovan; Pos.Sel.: Positively-selected regions.

| Gene | HS SNV | Nean/Den SNV | Pos.Sel. | rsID | Predicted microRNA |
| --- | --- | --- | --- | --- | --- |
| ***E2F6*** | ✓ | ✓ | ✓ | rs148715349 | hsa-miR-1280/455-3p |
| ***GLI3*** | ✓ | - | ✓ | rs77197280 | hsa-miR-335-5p/339-3p/4729 |
| ***RUNX2*** | - | ✓ | - | rs188598788 | hsa-miR-3143/4511/4803 |
| ***SOST*** | ✓ | - | - | rs17886183 | hsa-miR-5583-3p |

We performed a luciferase assay to functionally assess the impact of the species-specific 3’UTR variants of the four genes presented above. Vectors carrying the *Homo sapiens* allele and the counterpart Neanderthal/Denisovan allele for each gene were compared on the basis of the ratio of *Photinus pyrali* and *Renilla reniformis* luciferase activities (see Figure 2). A statistically significant result was found when analyzing the *SOST* (*p =* 0.0019; Wilcoxon rank sum test) and *RUNX2* (*p =* 0.0189; Wilcoxon rank sum test) 3’UTR variant effects. The rest of comparisons (for *E2F6* and *GLI3*) did not pass the statistical threshold set at *p <* 0.05. Thus, both a *Homo sapiens*-derived and a Neanderthal/Denisovan-derived 3’UTR SNV cause a differential regulation in *SOST* and *RUNX2* transcripts, respectively, in a bone cell-line model.

**Figure 2:**
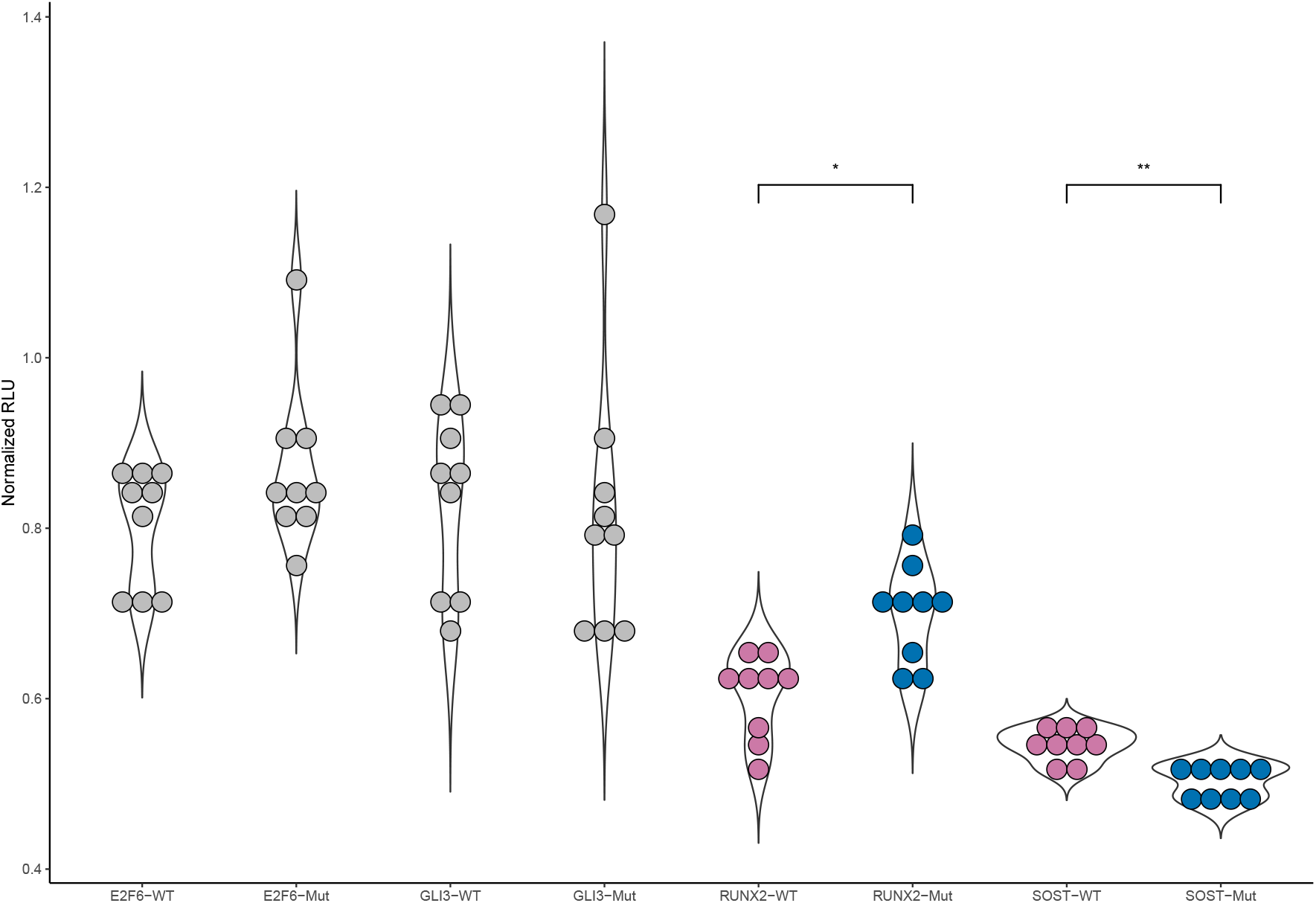
Luciferase reporter assay for the comparison *Homo sapiens versus*Neanderthal/Denisovan alleles. Normalized Relative Luciferase Units (RLU; Ratio of *Photinus pyrali* and *Renilla reniformis* luciferase activities) were compared between the *Homo sapiens* (WT) and Neanderthal/Denisovan variants (Mut) for each gene, applying a Wilcoxon rank sum test. Values are normalized to the activity of the empty vector, arbitrarily set at 1. Significant differences were found for *SOST* (*p*-value 0.0019) and *RUNX2* (*p*-value 0.0189) respective variant comparisons. No significant results were found for*E2F6* (*p*-value 0.3865) and *GLI3* (*p*-value 0.4363) *: *p <* 0.05; **: *p <* 0.01

## 3 Discussion

Inquiry into the ‘human condition’ is now being illuminated with the retrieval and analyses of ancient DNA material [9]. In this work, we investigated the role of *Homo*-specific genetic variants on transcript regulation. We selected four candidate genes known to affect skeletal and craniofacial phenotypes and whose 3’UTRs harbor derived variants in either the *Homo sapiens* lineage (*E2F6*, *GLI3* and *SOST*) or the Neanderthal/Denisovan lineage (*RUNX2*). The regulatory impact of each variant was assessed with a luciferase reporter assay in a bone cell-line model. We found that the *SOST* and *RUNX2* 3’UTR species-specific variants cause a differential regulation in the reporter expression assay, while no effect was detected for *E2F6* and *GLI3* 3’UTR variants.

The two Neanderthal/Denisovan 3’UTR variants identified affecting reporter expression above the threshold of statistical significance present distinct directional changes: A downregulation of *SOST* and an upregulation of *RUNX2* transcriptional activities. The assessment of the impact of these regulatory variants on recent human evolution is illuminated by the clinical conditions associated to the malfunction of the relevant genes. SOST acts as an inhibitor of the Wnt pathway that affects bone formation [40, 41] and, clinically, its deficiency has been associated with rare autosomal recessive bone dysplasias: Sclerosteosis, linked to loss-of-function mutations in sclerostin (encoded by *SOST*), and van Buchen disease, caused by a deletion of a region 35 kilobases down-stream of *SOST* affecting a regulatory element [36]. It has also been reported in patients with craniodiaphyseal dysplasia carrying mutations in the secretion peptide of sclerostin [37]. Patients with these diseases show excessive bone mass with specific facial morphologies, most prominently, forehead bossing and mandibular overgrowth [36]. Pronounced differences in bone density are observed when craniofacial features between *Homo sapiens* and Neanderthals are compared, with the former comparatively possessing reduced brow ridges, nasal projection and prognathism [21].

Of note, the *SOST* ancestral variant studied here has previously been found in five cases from a cohort of postmenopausal women with extreme values of BMD, four times in the high bone mass subgroup and one in the low bone mass subgroup in [42], as well as in a woman with high BMD in [43]. In accordance with these reports, we observed in an exploratory analysis that the mean value of BMD found in 30 individuals from a cohort of postmenopausal women (BARCOS cohort; [42]) carrying the heterozygous mutation rs17886183 is higher than in those carrying the reference allele (0.889 vs 0.850 g/cm^2^). The significance of this result will have to be determined in larger sample sizes because of the low frequency of this variant. Still, the directional change caused by the Neanderthal/Denisovan 3’UTR variant studied here is in agreement with the expected morphological effects (higher BMD) derived from lower levels of the SOST protein.

Turning to *RUNX2*, the gene is a critical regulator of skeletal development promoting osteoblast differentiation and chondrocyte maturation [44]. Loss-of-function mutations in the gene have been found in patients with cleidocranial dysplasia, an autosomal dominant disorder clinically characterized by short stature, incomplete and delayed closure of fontanelles, frontal bossing, hypoplasia of clavicles or bell shaped thorax [35]. As previously highlighted by [45, 46], some of the anatomical structures affected by *RUNX2* dysfunction are known to differ between our species and the Neanderthals: the closure of fontanelles may be delayed in *Homo sapiens* in light of the globularization phase that takes place after birth and only attributed to our species [30, 47–49], or the wider lower thorax in a characteristically bell shape present in the Neanderthals [2].

In the case of *RUNX2*, inferences about the phenotypic consequences of the differential protein dosage as the one observed in our experimental setting are more limited, given that studies in mice have shown that *Runx2* knockout mice completely lack ossification [50, 51], and *Runx2* overexpression causes an osteopenic phenotype [52]. Nevertheless, two association studies have reported a SNV (rs7771980) in one of the two promoters of *RUNX2*, where the nucleotide version present at high frequency in modern human populations is linked to lower BMD ([53, 54]; the homozygous rare version was reported to be associated with lower BMD in [55]). In addition, the promoter carrying this SNV derived in *Homo sapiens* (and other polymorphisms in linkage disequilibrium) shows lower activity than the counterpart carrying the alternative alleles on a reporter assay in a osteoblast-like rat cell-line [53]. In consonance with these findings, the *Homo sapiens*-derived 3’UTR tested in this study also causes lower *RUNX2* transcriptional activity.

In summary, the results obtained under this experimental model demonstrate *Homo* divergent regulation where the Neanderthal/Denisovan noncoding variants increase *RUNX2* and decrease *SOST* expression levels, underscoring the importance of 3’UTR variants as likely contributors to the phenotypic differences in bone mineral density between our species and our closest extinct relatives.

## 4 Methods

### 4.1 3’UTRs with species-specific changes

A dataset containing a catalog of SNVs derived in the AMH lineage at fixed or nearly fixed frequency-where the Neanderthals and the Denisovans carry the ancestral allele-, as well as alleles derived in the Neanderthal/Denisovan lineage [10], was processed and integrated with the Ensembl human assembly GRCh37 [27] to identify genes with: 1. 3’UTRs that contain derived SNVs in either the modern lineage or the Neanderthal/Denisovan lineage; 2. 3’UTRs that accumulated derived changes exclusively either in the modern human lineage or in the Neanderthal/Denisovan lineage. For Table 1: Coordinates of putative positively-selected regions for the intersection with 3’0UTR variants were retrieved from [26]. Likewise, microRNAs-SNP predictions were retrieved via [56].

#### Enrichment analysis

Genes with species-specific changes in their 3’UTRs were used for functional enrichment analysis using the *gProfiler2* R package [28], using the hypergeometric test with multiple comparison correction (method “gSCS”) and significance threshold set at p < 0.05.

### 4.2 Luciferase assay

#### Cell culture

The human osteosarcoma cell-line Saos-2 was used for luciferase reporter assays. It was obtained from the American Type Culture Collection (ATCC® HTB-85™) and grown in Dulbecco’s Modified Eagle Medium (DMEM; Sigma-Aldrich), supplemented with 10% Fetal Bovine Serum (FBS; Gibco, Life Technologies) and 1% penicillin/streptomycin (p/s; Gibco, Life Technologies), at 37°C and 5% of CO_2_.

#### Luciferase reporter constructs and site-directed mutagenesis

Fragments (around 400 bp in length) of the *SOST*, *E2F6*, *RUNX2* and *GLI3* 3’UTRs containing the SNPs rs17886183, rs148715349, rs144321470 and rs188598788, respectively, in a central position, were cloned (in each of the two allele versions) into the pmirGLO Dual-Luciferase miRNA target expression vector (Promega). Constructs were cloned using the XhoI and SdaI restriction sites. All primers used are detailed in Suppl. Table 1. In all cases, the presence of point mutations and absence of errors were verified through Sanger sequencing.

#### Reporter assay

1.1*x*10^5^ Saos-2 cells per well were cultured in 12-well plates, 24h before the transfection. We transfected at a 1:1 ratio, the pGFP vector and a vector containing either the empty pmirGLO or the *SOST*-3’UTR, *E2F6* -3’UTR, *RUNX2* -3’UTR or *GLI3* -3’UTR fragments (with either the *Homo sapiens* or the Neaderthal/Denisovan variants). Fugene HD was used following the manufacturers instructions. Forty-eight hours after transfection, cells were lysed and the luciferase activities of *Photinus pyralis* and *Renilla reniformis* were measured using a GloMax®-Multi luminometer (Promega), following the instructions of the Dual-luciferase reporter assay system (Promega). Each allelic variant was tested in triplicate and each experiment was repeated three times. Mean values for each variant (n = 9; normalized respect to empty vector) were used for statistical comparisons.

#### Statistics

Statistical significance for luciferase activity differences was evaluated applying the Wilcoxon rank sum test for each independent allelic comparison. Results were considered significant if p < 0.05.

#### *SOST* 3’UTR SNP genotyping

The *SOST* 3’UTR SNP (rs17886183) modifies the restriction enzyme sequence of NlaIII. We took advantage of it to genotype this SNP in 777 women of the postmenopausal BARCOS cohort [42]. For digestion with NlaIII, the PCR fragment was amplified with the primers detailed in Supplementary Table 1 and purified in MultiScreen TM Vacuum Manifold 96-well plates (Merck Millipore). The purified PCR fragment was incubated with either the enzyme NlaIII or water (negative control) for 4 h at 37 °C. Then we inactivated the enzyme by heating in the dry bath (65°C, 20 minutes). The digested samples were run on a 2% agarose gel (80V, 1 hour). 10% of the samples were sequenced by the Sanger method (CCiTUB genomic service, Parc Cientific, Barcelona, Spain) for genotyping quality control, and they showed 100% concordance. The tagging kit used is BigDye™ Terminator v3.1 Cycle Sequencing Kit, detection and electrophoresis were performed on automated capillary sequencer models 3730 Genetic Analyzer and 3730xl Genetic Analyzer. The dataset with BMD values and genotype from the BARCOS cohort can be found at https://github.com/jjaa-mp/3utr_humanevolution.

## Data and material availability

Datasets generated in this manuscript and code used for the analysis and generation of figures are made available through Supplementary Table 1 and at https://github.com/jjaa-mp/3utr_humanevolution.

## Acknowledgments

We thank Monica Cozar, Nicole Stender and Nerea Ugartondo for technical assistance.

## Author Contributions

Conceptualization: CB & DG & SB & AA & NMG & JM; Data Curation: AA & NMG & JM; Performance of experiments: NMG & JM; Formal Analysis: AA & NMG & JM; Visualization: AA & NMG & JM; Writing - Original Draft Preparation: CB & DG & SB & AA & NMG & JM; Writing - Review & Editing: CB & DG & SB & AA & NMG & JM; Supervision: CB & DG & SB; Funding Acquisition: CB.

## Funding statement

CB, SB, DG and NMG acknowledge support from the Spanish Ministry of Economy and Competitiveness (grants PID2019-107042GB-I00, SAF2016-75948-R and PID2019-107188RB-C2). CB also acknowledges support from the MEXT/JSPS Grant-in-Aid for Scientific Research on Innovative Areas #4903 (Evolinguistics: JP17H06379), and Generalitat de Catalunya (2017-SGR-341). JM acknowledges financial support from the Departament d’Empresa i Coneixement, Generalitat de Catalunya (FI-SDUR 2020). AA acknowledges financial support from the Spanish Ministry of Economy and Competitiveness and the European Social Fund (BES-2017-080366).

## Competing interests

The authors declare no competing interests.

## Notes

https://github.com/jjaa-mp/3utr_humanevolution

### Competing Interest Statement

The authors have declared no competing interest.

## References

[1] M. Bastir, et al., “Differential Growth and Development of the Upper and Lower Human Thorax,” PLOS ONE, vol. 8, p. e75128, Sept. 2013.

[2] D. Garía-Martínez, et al., “Early development of the Neanderthal ribcage reveals a different body shape at birth compared to modern humans,” Science Advances, vol. 6, p. eabb4377, Oct. 2020.

[3] A. Gómez-Olivencia, et al., “3D virtual reconstruction of the Kebara 2 Neandertal thorax,” Nature Communications, vol. 9, p. 4387, Oct. 2018.

[4] N. J. Friedlander et al., “Obstetric implications of neanderthal robusticity and bone density,” Human Evolution, vol. 9, pp. 331–342, Oct. 1994.

[5] J. D. Polk, “Influences of limb proportions and body size on locomotor kinematics in terrestrial primates and fossil hominins,” Journal of Human Evolution, vol. 47, pp. 237–252, Oct. 2004.

[6] M. Meyer, et al., “A High-Coverage Genome Sequence from an Archaic Denisovan Individual,” Science, vol. 338, pp. 222–226, Oct. 2012. Publisher: American Association for the Advancement of Science Section: Research Article.

[7] K. Prüfer, et al., “The complete genome sequence of a Neanderthal from the Altai Mountains,” Nature, vol. 505, pp. 43–49, Jan. 2014. Number: 7481 Publisher: Nature Publishing Group.

[8] K. Prüfer, et al., “A high-coverage Neandertal genome from Vindija Cave in Croatia,” Science, vol. 358, pp. 655–658, Nov. 2017. Publisher: American Association for the Advancement of Science Section: Report.

[9] S. Pääbo, “The Human Condition—A Molecular Approach,” Cell, vol. 157, pp. 216–226, Mar. 2014.

[10] M. Kuhlwilm et al., “A catalog of single nucleotide changes distinguishing modern humans from archaic hominins,” Scientific Reports, vol. 9, p. 8463, June 2019.

[11] S. Weyer et al., “Functional Analyses of Transcription Factor Binding Sites that Differ between Present-Day and Archaic Humans,” Molecular Biology and Evolution, vol. 33, pp. 316–322, Feb. 2016.

[12] T. Maricic, et al., “A Recent Evolutionary Change Affects a Regulatory Element in the Human FOXP2 Gene,” Molecular Biology and Evolution, vol. 30, pp. 844–852, Apr. 2013.

[13] H. R. Barker, et al., “Evolution is in the details: Regulatory differences in modern human and Neanderthal,” bioRxiv, p. 2020.09.04.282749, Feb. 2021.

[14] S. M. Yan et al., “Archaic hominin genomics provides a window into gene expression evolution,” Current Opinion in Genetics & Development, vol. 62, pp. 44–49, June 2020.

[15] A. W. Briggs, et al., “Removal of deaminated cytosines and detection of in vivo methylation in ancient DNA,” Nucleic Acids Research, vol. 38, pp. e87–e87, Apr. 2010.

[16] J. S. Pedersen, et al., “Genome-wide nucleosome map and cytosine methylation levels of an ancient human genome,” Genome Research, vol. 24, pp. 454–466, Mar. 2014.

[17] D. Gokhman, et al., “Reconstructing the DNA Methylation Maps of the Neandertal and the Denisovan,” Science, vol. 344, pp. 523–527, May 2014.

[18] D. Gokhman, et al., “Differential DNA methylation of vocal and facial anatomy genes in modern humans,” Nature Communications, vol. 11, p. 1189, Mar. 2020.

[19] D. Batyrev, et al., “Predicted Archaic 3D Genome Organization Reveals Genes Related to Head and Spinal Cord Separating Modern from Archaic Humans,” Cells, vol. 9, p. 48, Dec. 2019.

[20] D. Gokhman, et al., “Reconstructing Denisovan Anatomy Using DNA Methylation Maps,” Cell, vol. 179, pp. 180–192.e10, Sept. 2019.

[21] C. Theofanopoulou, et al., “Self-domestication in Homo sapiens: Insights from comparative genomics,” PLOS ONE, vol. 12, p. e0185306, Oct. 2017.

[22] M. Zanella, et al., “Dosage analysis of the 7q11.23 Williams region identifies BAZ1B as a major human gene patterning the modern human face and underlying self-domestication,” Science Advances, vol. 5, p. eaaw7908, Dec. 2019.

[23] C. V. Weiss, et al., “The cis-regulatory effects of modern human-specific variants,” preprint, Evolutionary Biology, Oct. 2020.

[24] K. A. Wanke, et al., “Understanding Neurodevelopmental Disorders: The Promise of Regulatory Variation in the 3UTRome,” Biological Psychiatry, vol. 83, pp. 548–557, Apr. 2018.

[25] M. Lopez-Valenzuela, et al., “An Ancestral miR-1304 Allele Present in Neanderthals Regulates Genes Involved in Enamel Formation and Could Explain Dental Differences with Modern Humans,” Molecular Biology and Evolution, vol. 29, pp. 1797–1806, July 2012.

[26] S. Peyrégne, et al., “Detecting ancient positive selection in humans using extended ineage sorting,” Genome Research, vol. 27, pp. 1563–1572, Sept. 2017.

[27] A. D. Yates, et al., “Ensembl 2020,” Nucleic Acids Research, vol. 48, pp. D682–D688, Jan. 2020.

[28] U. Raudvere, et al., “g:Profiler: a web server for functional enrichment analysis and conversions of gene lists (2019 update),” Nucleic Acids Research, vol. 47, pp. W191–W198, July 2019.

[29] J.-J. Hublin, et al., “Brain ontogeny and life history in Pleistocene hominins,” Philosophical Transactions of the Royal Society B: Biological Sciences, vol. 370, p. 20140062, Mar. 2015. Publisher: Royal Society.

[30] S. Neubauer, et al., “The evolution of modern human brain shape,” Science Advances, vol. 4, p. eaao5961, Jan. 2018. Publisher: American Association for the Advancement of Science Section: Research Article.

[31] T. Kochiyama, et al., “Reconstructing the Neanderthal brain using computational anatomy,” Scientific Reports, vol. 8, p. 6296, Apr. 2018. Number: 1 Publisher: Nature Publishing Group.

[32] K. Estrada, et al., “Genome-wide meta-analysis identifies 56 bone mineral density loci and reveals 14 loci associated with risk of fracture,” Nature Genetics, vol. 44, pp. 491–501, Apr. 2012.

[33] J. A. Morris, et al., “An atlas of genetic influences on osteoporosis in humans and mice,” Nature genetics, vol. 51, pp. 258–266, Feb. 2019.

[34] M. Kalff-Suske, et al., “Point mutations throughout the GLI3 gene cause Greig cephalopolysyndactyly syndrome,” Human Molecular Genetics, vol. 8, pp. 1769–1777, Sept. 1999.

[35] S. Mundlos, “Cleidocranial dysplasia: clinical and molecular genetics,” Journal of Medical Genetics, vol. 36, pp. 177–182, Mar. 1999.

[36] A. H. van Lierop, et al., “Sclerostin deficiency in humans,” Bone, vol. 96, pp. 51–62, Mar. 2017.

[37] S. J. Kim, et al., “Identification of signal peptide domain SOST mutations in autosomal dominant craniodiaphyseal dysplasia,” Human Genetics, vol. 129, pp. 497–502, May 2011.

[38] J. Storre, et al., “Homeotic transformations of the axial skeleton that accompany a targeted deletion of E2f6,” EMBO reports, vol. 3, pp. 695–700, July 2002.

[39] M. Courel, et al., “E2f6 and Bmi1 cooperate in axial skeletal development,” Developmental Dynamics: An Official Publication of the American Association of Anatomists, vol. 237, pp. 1232–1242, May 2008.

[40] L. I. Plotkin et al., “Osteocytic signalling pathways as therapeutic targets for bone fragility,” Nature Reviews Endocrinology, vol. 12, pp. 593–605, Oct. 2016.

[41] J. Delgado-Calle, et al., “Role and mechanism of action of sclerostin in bone,” Bone, vol. 96, pp. 29–37, Mar. 2017.

[42] N. Martínez-Gil, et al., “Common and rare variants of WNT16, DKK1 and SOST and their relationship with bone mineral density,” Scientific Reports, vol. 8, p. 10951, Dec. 2018.

[43] N. Martínez-Gil, et al., “Genetics and Genomics of SOST: Functional Analysis of Variants and Genomic Regulation in Osteoblasts,” International Journal of Molecular Sciences, vol. 22, p. 489, Jan. 2021. Number: 2 Publisher: Multidisciplinary Digital Publishing Institute.

[44] T. Komori, “Roles of Runx2 in Skeletal Development,” Advances in Experimental Medicine and Biology, vol. 962, pp. 83–93, 2017.

[45] R. E. Green, et al., “A Draft Sequence of the Neandertal Genome,” Science, vol. 328, pp. 710–722, May 2010. Publisher: American Association for the Advancement of Science Section: Research Article.

[46] M. Kuhlwilm, et al., “Identification of Putative Target Genes of the Transcription Factor RUNX2,” PLOS ONE, vol. 8, p. e83218, Dec. 2013. Publisher: Public Library of Science.

[47] S. Neubauer, et al., “Endocranial shape changes during growth in chimpanzees and humans: a morphometric analysis of unique and shared aspects,” Journal of Human Evolution, vol. 59, pp. 555–566, Nov. 2010.

[48] P. Gunz, et al., “Brain development after birth differs between Neanderthals and modern humans,” Current biology: CB, vol. 20, pp. R921–922, Nov. 2010.

[49] P. Gunz, et al., “A uniquely modern human pattern of endocranial development. Insights from a new cranial reconstruction of the Neandertal newborn from Mezmaiskaya,” Journal of Human Evolution, vol. 62, pp. 300–313, Feb. 2012.

[50] T. Komori, et al., “Targeted disruption of Cbfa1 results in a complete lack of bone formation owing to maturational arrest of osteoblasts,” Cell, vol. 89, pp. 755–764, May 1997.

[51] F. Otto, et al., “Cbfa1, a candidate gene for cleidocranial dysplasia syndrome, is essential for osteoblast differentiation and bone development,” Cell, vol. 89, pp. 765–771, May 1997.

[52] W. Liu, et al., “Overexpression of Cbfa1 in osteoblasts inhibits osteoblast maturation and causes osteopenia with multiple fractures,” The Journal of Cell Biology, vol. 155, pp. 157–166, Oct. 2001.

[53] J. D. Doecke, et al., “Association of functionally different RUNX2 P2 promoter alleles with BMD,” Journal of Bone and Mineral Research: The Official Journal of the American Society for Bone and Mineral Research, vol. 21, pp. 265–273, Feb. 2006.

[54] M. Bustamante, et al., “Promoter 2 - 1025 T/C Polymorphism in the RUNX2 Gene Is Associated with Femoral Neck BMD in Spanish Postmenopausal Women,” Calcified Tissue International, vol. 81, pp. 327–332, Oct. 2007.

[55] H.-J. Lee, et al., “Association of a RUNX2 Promoter Polymorphism with Bone Mineral Density in Postmenopausal Korean Women,” Calcified Tissue International, vol. 84, pp. 439–445, June 2009.

[56] C. Liu, et al., “MirSNP, a database of polymorphisms altering miRNA target sites, identifies miRNA-related SNPs in GWAS SNPs and eQTLs,” BMC Genomics, vol. 13, p. 661, Nov. 2012.

